# Combined biosynthesis and site-specific incorporation of phenylalanine derivatives from aryl aldehydes or carboxylic acids in engineered bacteria

**DOI:** 10.1101/2024.12.17.628963

**Authors:** Shelby R. Anderson, Abigail P. Spangler, Michaela A. Jones, Roman M. Dickey, Aditya M. Kunjapur

## Abstract

Applications of genetic code expansion in live cells are widespread and continually emerging, yet they have been limited by their reliance on the supplementation of non-standard amino acids (nsAAs) to cell culturing media. While advances in cell-free biocatalysis are improving nsAA synthesis cost and sustainability, such processes remain reliant on multi-step processes of product isolation followed by supplementation to engineered cells. Here, we report the design of a modular and genetically encoded system that combines the steps of biosynthesis of diverse phenylalanine derivatives, which are the most frequently used family of nsAAs for genetic code expansion, and their site-specific incorporation within target proteins using a single engineered bacterial host. Unlike previous demonstrations in which individual nsAAs were targeted for biosynthesis and site-specific incorporation, our system serves as a platform that exhibits broad substrate specificity towards commercially ubiquitous, achiral building blocks of aryl aldehydes or carboxylic acids. We demonstrate that this modular system enables high fidelity biosynthesis and incorporation of nsAAs for multiple industrially relevant nsAAs, such as the incorporation of 4,4-L-biphenylalanine within proteins after supplementation with biphenylaldehyde and the incorporation of 4-azido-L-phenylalanine within proteins after supplementation with 4-azido-benzoic acid. Finally, we show that the combination of nsAA biosynthesis and incorporation steps can extend the chemical reach of the intrinsic biological containment strategy of synthetic auxotrophy from nsAAs to low-cost and achiral building blocks. We anticipate that our system will aid industrial-scale manufacturing of proteins that contain nsAAs and democratize access to expensive or commercially unavailable chemistries for labs that lack separations or synthesis expertise.

## Introduction

Genetic code expansion enables chemical diversification of proteins in living systems for a growing number of applications^1^, including the manufacture of conjugation-ready therapeutic proteins^2^ and intrinsic biological containment of engineered microbes^3^. Many of these applications are enabled by site-specific ribosomal incorporation of non-standard amino acids (nsAAs) that are derivatives of phenylalanine, such as 4-azido-L-phenylalanine, 4-acetyl-L-phenylalanine, and 4,4-L-biphenylalanine. Currently, these and other valuable phenylalanine derivatives are produced by chemical synthesis^4^, though several promising biocatalytic or photobiocatalytic synthesis platforms are under development by ourselves and others^5–10^. However, whether nsAA synthesis occurs chemically or biologically, the current synthesis paradigms for genetic code expansion applications necessitate downstream processing steps of nsAA isolation and supplementation to cell culture media. In addition to the costs and labor of the nsAA synthesis and isolation steps, the requirement of nsAA supplementation also imposes constraints that limit the performance or scalability of genetic code expansion in diverse application contexts.^11^ For example, in bioreactor settings, which may be highly relevant for on-site manufacturing of therapeutics, these constraints can include the cost and cellular uptake rate of the nsAA. For emerging contexts in which engineered microbes possess intrinsic biocontainment systems for controlled release in open environments, these can include the requirement that conditions permissive for growth must contain a specific nsAA.

Combined nsAA biosynthesis and site-specific incorporation within proteins using a single engineered organism could address these limitations and create opportunities to harness principles from synthetic biology, such as on-demand nsAA synthesis, nsAA sensing, and intercellular communication. For these and other reasons, pioneering efforts during the last two decades have begun to demonstrate coupling of nsAA biosynthesis and site-specific incorporation for certain nsAAs.^11–23^ However, past efforts featured biosynthetic pathways that exhibit highly restricted substrate specificity such that each desired nsAA requires a different pathway. Recent efforts have begun to show that closely related halogenated tyrosine^24^ and tryptophan^25^ analogs can be biosynthesized and incorporated into proteins; however, the breadth of substrates is still limited in their documented application space thus far. In contrast, biocatalysis-oriented efforts have leveraged polyspecific enzymes as platforms for synthesis of diverse phenylalanine derivatives in cell-free contexts. However, these biocatalytic strategies have not been demonstrated to function in the context of growing cells that are most actively translating new proteins, which is necessary when combining biosynthesis and incorporation.

Given the extensive application of >80 phenylalanine derivatives in genetic code expansion technologies and the lack of a promiscuous biosynthetic pathway to access these encodable nsAAs, we aimed to design a platform to produce these chemistries from inexpensive and readily accessible precursors. We imposed four requirements on our envisioned biosynthetic pathway design: (1) accept inexpensive and commercially available substrates; (2) employ promiscuous enzymes to access diverse nsAA targets from diverse substrates; (3) produce the desired nsAA in fermentative contexts; and, (4) function with high-fidelity when co-expressed with an orthogonal translation system (OTS) for combined nsAA biosynthesis and site-specific incorporation.

In relation to the first two requirements, we^26^ and others^27^ recently demonstrated the design of efficient and polyspecific three-enzyme cell-free biocatalytic cascades for synthesis of phenylalanine derivatives from inexpensive aryl aldehyde precursors. Our cascade features an L-Threonine Transaldolase (L-TTA) from *Pseudomonas fluorescens* (ObiH)^28–31^, a phenylserine dehydratase from *Ralstonia piketti* (*Rp*PSDH)^32^, and an aromatic aminotransferase from *E. coli* (TyrB). The cascade proceeds through two intermediates - a β-hydroxylated α-amino acid and a keto acid - both of which are high-value products on their own.^33^ Additionally, we showed that a carboxylic acid reductase (CAR) from *Segniliparus rotundus* (*Sr*CAR) could be added to the start of the cascade to use aryl carboxylic acids as starting materials, which may be less expensive, less volatile, less reactive, and more soluble than aldehydes.

Inspired by the success of this cascade in dilute aqueous settings, here we investigate what it would take to perform similar reactions in live cells, which can pose several potential challenges as well as the opportunity for process intensification. Using strains of *E. coli* that are engineered to stabilize aryl aldehydes,^34–36^ we show the successful biosynthesis of multiple highly relevant nsAAs that vary in size and polarity from either aryl aldehyde or carboxylic acid precursors in metabolically active cells. We proceed to demonstrate the coupling of biosynthesis and high-fidelity encoding of multiple nsAAs starting from either an aldehyde or carboxylic acid precursor. Finally, we illustrate how this platform for bioconversion from readily accessible precursors to nsAAs offers new synthetic biology applications based on control of protein translation by provision of aryl aldehydes, such as extending the permissive chemical range of the biological containment technique of synthetic auxotrophy to synthetic aryl aldehydes.

## Materials and Methods

### Strains and plasmids

Molecular cloning and vector propagation were performed in DH5α. Polymerase chain reaction (PCR) based DNA replication was performed using KOD XTREME Hot Start Polymerase. Cloning was performed using Gibson Assembly. Oligos were purchased from Integrated DNA Technologies (IDT).

### Materials and Chemicals

The following compounds were purchased from MilliporeSigma: Kanamycin sulfate (Kan), chloramphenicol (Cm), streptomycin sulfate salt (Str), dimethyl sulfoxide (DMSO), potassium phosphate dibasic, potassium phosphate monobasic, magnesium sulfate, calcium chloride dihydrate, glycerol, Tris base, glycine, HEPES, ATP, and KOD XTREME Hot Start polymerase. The following were purchased from TCI America: D-glucose, phenylpyruvic acid, biphenyl-4-carboxaldehyde. The following were purchased from Peptech: 4-acetyl-L-phenylalanine, 4-cyano-L-phenylalanine. The following were purchased from Sigma-Aldrich: benzaldehyde, biphenyl-4-carboxylic acid, L-phenylalanine. Agarose, ethanol, L-glutamic acid monopotassium salt monohydrate, and L-threonine were purchased from Alfa Aesar. Acetonitrile, trifluoroacetic acid (TFA), sodium chloride, LB Broth powder (Lennox), and LB Agar powder (Lennox) were purchased from Fisher Chemical. Taq DNA ligase was purchased from GoldBio. L-glutamic acid, was purchased from ACROS Organics. 4-cyanobenzoic acid and 4-azidobenzaldehyde were purchased from ChemCruz. 4-azido-L-phenylalanine was purchased from Bachem. Biphenylalanine was purchased from Combi-blocks. Phusion DNA polymerase and T5 exonuclease were purchased from New England BioLabs (NEB). Sybr Safe DNA gel stain was purchased from Invitrogen. NADPH (tetrasodium salt) and 4-acetylbenzaldehyde were purchased from Santa Cruz Biotechnology. Anhydrotetracycline (aTc) was purchased from Cayman Chemical. KAPA2G FAST Multiplex Kit was purchased from Roche.

### Culture Conditions

Cultures were grown in LB-Lennox medium (LBL: 10 g/L bacto tryptone, 5 g/L sodium chloride, 5 g/L yeast extract), or MOPS EZ Rich defined media (Teknova M2105) with 2% glucose unless otherwise specified.

### Stability Assays

For metabolically active cell stability testing, cultures of each *E. coli* strain to be tested were inoculated from a frozen glycerol stock (stored at -80 °C) and grown overnight in 3-5 mL of LBL media. Overnight cultures were then used to inoculate experimental cultures in 300 µL volumes in a 96-deep-well plate (Thermo Scientific™ 260251) at 100x dilution. Cultures were supplemented with 1 mM of aldehyde substrates (prepared in 100 mM stocks in DMSO) at mid-exponential phase (OD_600_∼0.5). Cultures were incubated at 37 °C with shaking at 1000 RPM and an orbital radius of 3 mm. Samples were taken by pipetting 200 µL from the cultures, centrifuging 4000 × *g* at 4 °C in a round bottom plate (SPL Life Sciences ISO 13485) and collecting the extracellular supernatant. Compounds were quantified after 20 h period using HPLC.

### Biosynthesis Assays for CAR and TTA screening

To prepare the strains for biosynthesis assays, plasmids were transformed into the RARE.Δ16 strain, unless otherwise specified. For the in vivo L-TTA activity assay, *E. coli* RARE.Δ16 was transformed with a pZE plasmid encoding expression of ObiH with a hexahistidine-SUMO tag at the N-terminus under the control of an aTc-inducible promoter. The strain was inoculated from frozen stocks and grown overnight in 3 mL LBL containing 30 µg/mL kanamycin (Kan). Overnight cultures were used to inoculate 300 µL volumes of MOPS EZ Rich media with 2% glucose and 100 mM L-Thr at a 1:100 inoculation ratio in a 96-deep-well plate (Thermo Scientific™ 260251) with appropriate antibiotic (30 µg/mL Kan). Cultures were incubated at 37 °C, shaking 1000 RPM with an orbital radius of 1.25 mm until and OD_600_ ∼ 0.5 was reached. OD_600_ was measured using a spectrophotometer. Then, 0.2 µM aTc was added for TTA expression and 1 mM of aldehyde precursor (**1b-4b**, prepared in 100 mM stocks in DMSO) was supplemented to the culture. Cultures were then incubated for 20 h at 30 °C. Samples were taken by pipetting 200 µL from the cultures, centrifuging 4000 × *g* at 4 °C in a round bottom plate (SPL Life Sciences ISO 13485) and collecting the supernatant. Metabolite concentration was measured via analysis of the supernatant on HPLC. For the L-TTA variant screening, RARE.Δ16 was transformed with a pZE plasmid encoding expression of *Pi*TTA, *Cs*TTA, *Bu*TTA, *Ka*TTA, or *Pb*TTA, all with a hexahistine-SUMO tag at the N-terminus. In vivo activity assays were performed as described above.

Cultures of each *E. coli* strain to be tested were inoculated from a frozen stock and grown overnight in 3 mL LB media containing 30 µg/mL Kan. Overnight cultures were then used to inoculate experimental cultures in 300 µL volumes in 96-deep-well plate (Thermo Scientific™ 260251) at 100x dilution and grown at 37 °C in MOPS EZ Rich media with 2 % glucose and Kan. At mid-exponential phase (OD_600_∼0.5), cultures were induced with 0.2 µM aTc and 2 mM of substrate (prepared in 100 mM stocks in DMSO) was added. Cultures were then incubated at 30 °C with shaking at 1000 RPM and an orbital radius of 3 mm. Samples were taken by pipetting 200 µL from the cultures, centrifuging in a round bottom plate (SPL Life Sciences ISO 13485) and collecting the extracellular broth. Compounds were quantified after a 4 h period using HPLC.

For the CAR variant screening, RARE.Δ16 was transformed with a pZE plasmid containing the Sfp gene from Bacillus subtilis and encoding aTc-inducible expression of *Sr*CAR, *Mav*CAR, *Mm*CAR, *Mab*CAR, *Ni*CAR, or *Af*CAR, all with a hexahistidine-tag at the N-terminus. In vivo activity assays were performed as described above, with metabolite concentration measured via HPLC after 2 h growth post induction and addition of 2 mM of the carboxylic acids **1a**-**3a** or 1 mM of **4a** (due to lower solubility).

### PSDH Expression and Activity Assays

For the in vivo PSDH assays, *E. coli* BL21 was transformed with a pZE plasmid encoding aTc-inducible expression of the *Rp*PSDH with a hexahistidine tag or a hexahistidine-SUMO tag at the N-terminus. Strains were inoculated from frozen stocks and grown overnight in 5 mL LBL containing 30 µg/mL Kan. Overnight cultures were used to inoculate 300 µL volumes of MOPS EZ Rich media with 0.4% glucose at a 1:100 inoculation ratio in a 96-deep-well plate with appropriate antibiotic (30 µg/mL Kan). Cultures were incubated at 37 °C, shaking 1000 RPM until and OD_600_ ∼ 0.5, as described previously, then 0.2 μM aTc was added for PSDH expression and 5 mM of substrate, L-*threo*-phenylserine, was supplemented to the culture. Cultures were then incubated for 20 h at 30 °C and supernatant collected for analysis via HPLC.

To evaluate *Rp*PSDH expression with and without the SUMO tag, overnight cultures were used to inoculate 3 mL LBL containing 30 µg/mL Kan and incubated at 37 °C in a shaking incubator at 250 RPM until an OD600 of 0.5-0.8 was reached. PSDH expression was induced by the addition of 0.2 µM aTc and cultures were incubated at 30 °C for 18 h. Then, 1 mL of cells was transferred to a microcentrifuge tube and lysed with 0.05 mL of glass beads by vortexing using a Vortex Genie 2 for 15-30 min. Samples were centrifuged at 18,213 × *g* at 4 °C for 15 min, lysate collected and subsequently denatured in Laemmli SDS reducing sample buffer for 10 min at 95 °C. Sample was then run on a sodium dodecyl-sulfate polyacrylamide gel electrophoresis (SDS-PAGE) gel with a Thermo Scientific^TM^ Spectra^TM^ Multicolor Broad Range Protein ladder and then analyzed via western blot with an HRP-conjugated 6*His, His-Tag Mouse McAB primary antibody. An Amersham ECL Primer chemiluminescent detection reagent was used to visualize the blot.

### Coupled TTA and PSDH Biosynthesis Assays

*E. coli* RARE.Δ16 was transformed with a plasmid with a colA origin of replication (pCola) encoding IPTG-inducible expression of both *Rp*PSDH with a hexahistidine-SUMO tag at the N-terminus and a TTA with a hexahistidine-SUMO tag at the N-terminus (either ObiH or *Pb*TTA). Overnight cultures were used to inoculate 3 mL LBL containing 50 µg/mL Carb. Overnight cultures were used to inoculate 300 µL volumes of MOPS EZ Rich media with 2% glucose and 100 mM L-Thr at a 1:100 inoculation ratio in a 96-deep-well plate with appropriate antibiotic (50 µg/mL Carb). Cultures were incubated at 37 °C in a shaking incubator at 1000 RPM. At mid-exponential phase (OD_600_∼0.5), cultures were induced with 1 mM IPTG and supplemented with 1 mM of aldehydes **1b-4b** (prepared in 100 mM stocks in DMSO) was added. Product concentration was quantified after 20 h incubation at 30 °C using HPLC.

### Coupled CAR, TTA and PSDH Biosynthesis Assays

*E. coli* RARE.Δ16 was transformed with two plasmids 1) a pCola vector encoding expression of both s-*Rp*PSDH and a L-TTA (either s-ObiH or s-*Pb*TTA) and 2) a pACYC vector encoding expression of *Mav*CAR and *Bs*Sfp. Overnight cultures were used to inoculate 3 mL LBL containing 25 µg/mL Carb and 17 µg/mL Cm. Overnight cultures were used to inoculate 300 µL volumes of MOPS EZ Rich media with 2% glucose and 100 mM L-Thr at a 1:100 inoculation ratio in a 96-deep-well plate with appropriate antibiotic. Cultures were incubated at 37 °C in a shaking incubator at 250 RPM. At mid-exponential phase (OD_600_∼0.5), cultures were induced with 1 mM IPTG and 1 mM of carboxylic acids **1a-4a** (prepared in 100 mM stocks in DMSO) was added. Product concentration was quantified after 20 h incubation at 30 °C using HPLC.

### Non-standard amino acid incorporation assays

*E. coli* RARE.Δ16 was transformed with pEVOL plasmid for arabinose-inducible expression of orthogonal AARS/tRNA pairs and a pZE plasmid for aTc-inducible expression of a reporter protein fusion consisting of a ubiquitin domain, followed by an in-frame amber suppression codon, followed by GFP (pZE-Ub-UAG-GFP). These strains were used to inoculate 300 µL volumes of MOPS EZ Rich media with 2% glycerol in a 96-deep-well plate with appropriate antibiotic 15 µg/mL Kan and 17 µg/mL Cm and were incubated at 37 °C in a shaking incubator at 1000 RPM. At mid-exponential phase (OD_600_ ∼0.5), cultures were induced with 0.2 µM aTc and 0.2% (wt/vol) L-arabinose and grown in the absence or presence of nsAA (supplemented at varying concentrations) for 18 h at 30 °C. Cells were then pelleted via centrifugation (4,000 x g at 4 °C for 12 min), washed in 1x PBS buffer, and then quantified for OD_600_ and GFP fluorescence at excitation and emission wavelengths of 485 and 525 nm, respectively, using a Spectramax i3x plate reader with Softmax Pro 7.0.3 software.

### Coupled biosynthesis and incorporation

For the conversion of externally supplemented aldehyde **4b** for subsequent incorporation of **4e**, *E. coli* RARE.Δ16 was co-transformed with 1) a single reporter/AARS/tRNA plasmid denoted pRepSynth-BipARS encoding expression of an ubiquitin-fused GFP reporter containing an amber suppression codon and a C-terminal hexahistidine tag and BipARS/tRNA^CUA^ and 2) a pCola vector encoding expression of both s-*Pb*TTA and s-*Rp*PSDH. This strain was cultured at 37 °C in 150 mL of LBL media with 100 mM L-Thr in a 500 mL baffled shake flask. At an OD_600_∼0.4, the culture was induced with 1 mM IPTG and 0.2 µM aTc and 2 mM 4b precursor was added. Cultures were then grown at 30 °C for an additional 18 h. The cells were harvested by centrifugation (4,000 × *g* at 4 °C for 5 min), supernatant was removed, and the pellet was stored at -80 °C pending purification.

For the conversion of externally supplemented aldehyde **1a** for subsequent incorporation of **1e**, *E. coli* ROAR^35^ was transformed with a three plasmid system consisting of 1) a single reporter/AARS/tRNA plasmid denoted pRepSynth-NapARS encoding aTc-inducible expression of an ubiquitin-fused GFP reporter containing an amber suppression codon and a C-terminal hexahistidine tag and NapARS/tRNA^CUA^, 2) a pCola vector encoding IPTG-inducible expression of both s-*Pb*TTA and s-*Rp*PSDH and 3) a pZE vector encoding aTc-inducible expression of *Mav*CAR and *Bs*Sfp. This strain was cultured at 37 °C in 75 mL of LBL media with 100 mM L-Thr in a 250 mL baffled shake flask in duplicate. At an OD_600_ ∼0.4, the culture was induced with 1 mM IPTG and 0.2 µM aTc and 2 mM **1a** precursor was added. Cultures were then grown at 30 °C for an additional 18 h. The cells were harvested by centrifugation, supernatant was removed, and the pellet was stored at -80 °C prior to purification.

For purification, the pellet was resuspended in 6-8 mL of lysis buffer (25 mM HEPES, 10 mM imidazole, 300 mM NaCl, pH 7.4) and disrupted using a QSonica Q125 sonicator at with cycles of 5 s at 90% amplitude and 10 s off for 8 min. The lysate was centrifuged in 1 mL aliquots in microcentrifuge tubes for 45 min at 18,213 × *g* at 4 °C. The supernatant was sterile filtered through a 0.22 µm syringe filter and purified using an Ni-Sepharose affinity chromatography (HisTrap HP, 5 mL) via an ÄKTA Pure GE fast protein liquid chromatography (FPLC) system using Lysis Buffer A (25 mM HEPES pH 7.4, 250 mM NaCl, 0.4 mM PLP, 10 mM MgCl_2_, and 10 mM imidazole) and Elution Buffer B (25 mM HEPES pH 7.4, 250 mM NaCl, 0.4 mM PLP, 10 mM MgCl_2_, and 250 mM imidazole). The column was equilibrated with 2.5% Buffer B and after performing an isocratic 12% Buffer B (40 mM) and 18% Buffer B (54 mM imidazole) wash, elution of the His-tagged proteins (reporter, TTA and PSDH, and CAR) was performed at 250 mM imidazole. Samples were subsequently denatured in Laemmli SDS reducing sample buffer for 10 min at 95 °C. Sample was then run on an SDS-PAGE gel with a Thermo Scientific^TM^ Spectra^TM^ Multicolor Broad Range Protein ladder to confirm reporter protein production and purification by size. The gel was then stained with Bio-Safe Coomassie stain (Bio-Rad), and destained with water.

For LC–MS/MS measurements, protein bands corresponding to the molecular weight of the protein of interest were cut from the polyacrylamide gel, and the gel fragments were digested by in-gel tryptic digestion. Extracted peptides were desalted using Pierce pipette tips (Thermo Fisher Scientific), dried, and prepared for LC–MS/MS using a Thermo Fisher Scientific Orbitrap Eclipse Tribrid Mass Spectrometer (MS; Thermo Fisher Scientific) with an Ultimate 3000 nano-LC and a FAIMS Pro Interface (Thermo Fisher Scientific) following the protocol detailed in Butler, et al.^13^ Proteomic analysis was performed in the MaxQuant-Andromeda software suite (v2.5.1.0). An *E. coli* reference proteome (taxonomy_id:83333) was used for the database search and the reporter protein with Tyr at the site for nsAA incorporation was used as the reference protein. Trypsin was specified at the digestion mode with 2 maximum missed cleavages. Variable modifications included methionine oxidation, Phe, Trp, **4e**, **1e**, or 4-amino-phenylalanine (pAF) substitutions for Tyr to account for potential misincorporation, reduction, or the target products. Other parameters were set to the default. Peptide intensity values from MaxQuant were used for quantification.

### Synthetic Auxotroph growth on aldehyde precursor

*E. coli* DEP.e5 was transformed with a pZE vector encoding aTc-inducible expression of both s-ObiH and s-*Rp*PSDH. Cultures were inoculated from a frozen stock and grown overnight in 3 mL LB media with 12.75 µg/mL Cm, 15 µg/mL Kan, 10 μM biphenylalanine (**4e**), 100 µM biphenylaldehyde (**4b**), 0.2% (wt/vol) L-arabinose, 0.2 µM aTc, 20 mM Tris-HCl pH 8, 0.005% SDS, and 100 mM L-Thr at 34 °C. Cells were washed 4x in LB media with 12.75 µg/mL Cm, 25 µg/mL Kan, 0.2% (wt/vol) L-arabinose, 0.2 µM aTc, 20 mM Tris-HCl pH 8, 0.005% SDS, and 100 mM L-Thr to remove the essential nutrient of biphenylalanine. Overnight cultures were then used to inoculate experimental cultures in 200 μL volumes a Greiner clear bottom 96 well plate (Greiner 655090) at 100x dilution in LB media with 12.75 µg/mL Cm, 15 µg/mL Kan, 0.2% (wt/vol) L-arabinose, 0.2 µM aTc, 20 mM Tris-HCl pH 8, 0.005% SDS, and 100 mM L-Thr. Substrate was added at inoculation under the following experimental conditions performed in triplicate: no substrate, 10 µM **4e**, or 100-500 µM aldehyde precursor **4b**. Cultures were grown for 50 h in a Spectramax i3x plate reader with medium plate shaking at 30 °C and with absorbance readings at 600 nm.

After 50 h growth, 5 µL of each experimental triplicate was plated on both non-permissive and permissive LB-agar to the confirm growth observed in liquid culture was due to the presence of supplemented or biosynthesized essential nutrient, rather than escape of the synthetic auxotroph. Non-permissive agar contained 12.75 µg/mL Cm, 15 µg/mL Kan, 0.2% (wt/vol) L-arabinose, 20 mM Tris-HCl pH 8, 0.005% SDS. Permissive agar included the essential nutrient biphenylalanine (**4e**) at 10 µM. Plates were incubated at 30 °C for 18 h.

### Liquid chromatography analysis

Metabolites and synthetic biochemicals of interest were quantified via high-performance liquid chromatography (HPLC) using an Agilent 1100 Infinity model equipped with a Zorbax Eclipse Plus-C18 column (part number: 959701-902, 5 µm, 95Å, 2.1 x 150 mm). To quantify compounds of interest, an initial mobile phase of solvent A/B = 95/5 was used (solvent A, water, 0.1% trifluoroacetic acid; solvent B, acetonitrile, 0.1% trifluoroacetic acid) and maintained for 5 min. A gradient elution was performed (A/B) with: gradient from 95/5 to 50/50 for 5-12 min, gradient from 50/50 to 0/100 for 12-13 min, gradient from 0/100 to 95/5 for 13-14, and equilibration at 95/5 for 14-15 min. A flow rate of 1 mL min^-1^ was maintained, and absorption was monitored at 210, 250, 270, 280 and 300 nm.

## Results

### Biosynthesis of model beta-hydroxylated nsAAs in live cells

We selected four model substrates that we envisioned could correspond to widely used nsAAs for applications such as bio-orthogonal conjugation, photo-crosslinking, and biological containment: 4-azido-benzaldehyde (**1b**) for biosynthesis of 4-azido-L-phenylalanine (**1e**); 4-acetyl-benzaldehyde (**2b**) for biosynthesis of 4-acetyl-L-phenylalanine (**2e**), 4-cyanobenzaldehyde (**3b**) for biosynthesis of 4-cyano-L-phenylalanine (**3e**), and 4,4-biphenylaldehyde (**4b**) for biosynthesis of 4,4-L-biphenylalanine (**4e**) (**Fig. 2a**). Besides their utility for applications, these structurally and electronically diverse phenylalanine derivatives could shed light on the compatibility of related chemistries in live-cell contexts. In our recent design of a one-pot biocatalytic cascade, we determined that a SUMO-tagged variant of the L-TTA from *Pseudomonas fluorescens*, s-ObiH, can accept these four model aldehyde substrates.^26^

**Figure 1.**
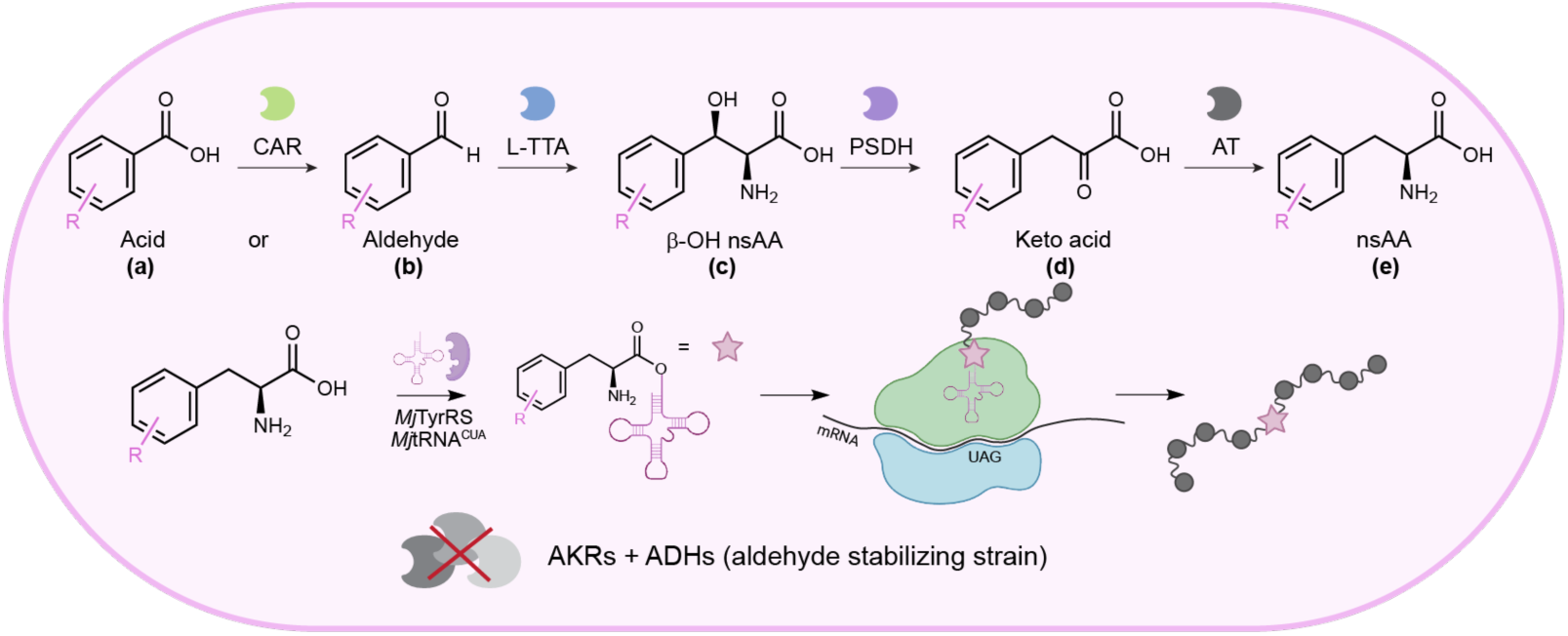
Overview of the engineered biosynthesis and incorporation platform. A single strain was designed to express two to four enzymes for the biosynthesis of phenylalanine derivates from supplemented acid or aldehyde precursors. The biosynthesized nsAA was then incorporated into a target protein using amber stop codon suppression and an orthogonal translation system.

**Figure 2.**
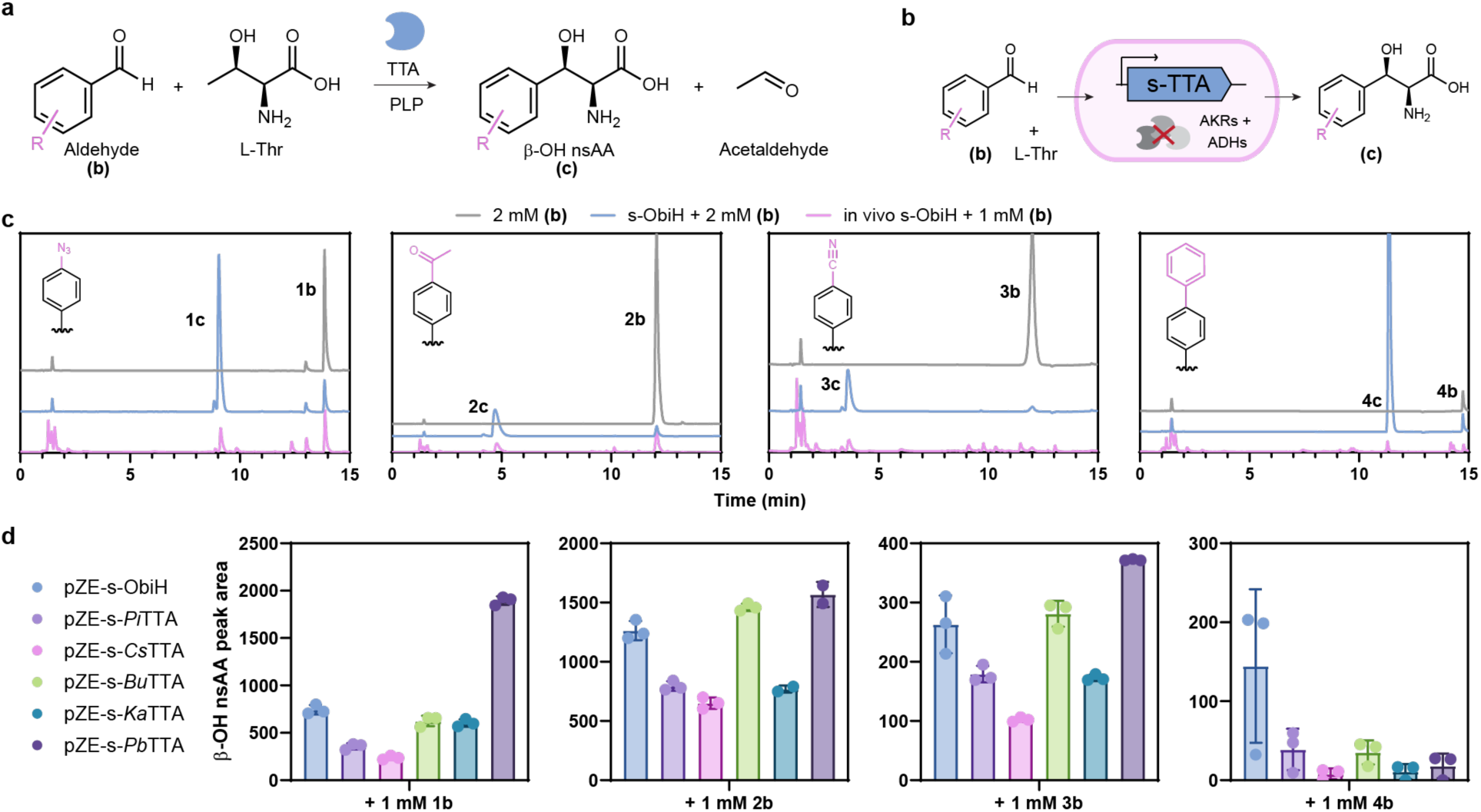
Reactions catalyzed by engineered live *E. coli* strains that were transformed for heterologous expression of L-threonine transaldolases (L-TTAs), which catalyze the first step of the pathway starting from aldehyde substrates. (A) Reaction catalyzed by L-TTA. (B) Illustration of the reaction format with a live strain engineered both for aldehyde retention and for expression of one of six possible SUMO-tagged L-TTAs. (C) HPLC traces obtained when supplying either just the aldehyde substrate (2 mM), an endpoint reaction catalyzed by the purified s-ObiH after provision of 2 mM aldehyde substrate (to determine the RT of the expected β-OH nsAA product), or an endpoint reaction catalyzed by cells that express s-ObiH and were supplied 1 mM aldehyde substrate. Four different aldehyde substrates were tested (#**b**), each corresponding to a commonly used nsAA for genetic code expansion. (D) Endpoint HPLC measurements demonstrating how the peak area of the expected β-OH nsAA product varies based on the L-TTA homolog that is chosen for heterologous expression in live cells.

We examined the rate of formation of β-hydroxylated nsAAs upon supplementing these aldehyde substrates at 1 mM concentrations to cultures of *E. coli* cells that were transformed for heterologous expression of s-ObiH (**Fig. 2b**). We chose a strain of *E. coli* whose genome we engineered for improved aldehyde stabilization. This strain, *E. coli* RARE.Δ16^36^, contains 6 gene knockouts from its progenitor *E. coli* RARE strain^34^ and 10 additional aldehyde reductase gene inactivations that enable the stabilization of the di-aldehyde terepthalaldehyde. We cultured the *E. coli* RARE.Δ16 strain in MOPS EZ Rich media with 2% glucose and 100 mM L-Thr. We were pleased to observe the desired β-hydroxylated nsAA products in the supernatant by HPLC at an endpoint of 20 h (**Fig. 2c**). Next, we screened s-ObiH and five SUMO-tagged homologs of ObiH with the model aldehydes as before to ascertain the role of L-TTA homolog selection on β-hydroxylated nsAA titer. All variants were active in live cell contexts under the conditions tested, with the optimal L-TTA being chemistry-specific (**Fig. 2d**). For the biosynthesis of **2c** and **3c**, s-*Pb*TTA was the best performer, though peak areas were comparable between it, s-*Bu*TTA, and s-ObiH. In contrast, for the biosynthesis of **4c**, s-ObiH generated over two-fold the peak area of any other L-TTAs tested, with two replicates resulting in over four-fold compared to the next best strain.

### Establishing a full biosynthetic pathway from aldehydes to nsAAs in live cells

We briefly evaluated the potential for site-specific incorporation of the β-hydroxylated nsAA to determine if sidechain chemistries of interest could be introduced within proteins without need for removal of the β-hydroxy group. However, screening via a routine nsAA incorporation assay with a commercially available β-hydroxylated nsAA (β-OH *O-*methyl-L-tyrosine) indicated that it was not a substrate for a large panel of published MjTyrRS/tRNA_CUA_ pairs. Accordingly, we gained additional rationale for β-hydroxy group removal and sought to accomplish this by extension of the pathway to include a phenylserine dehydratase from *Ralstonia piketti* (*Rp*PSDH) and the native aromatic aminotransferase in *E. coli* (TyrB) (**Fig. 3a**). Given the improvement of protein yield we observed from addition of a SUMO-tag to diverse L-TTAs, we inquired about its effect on *Rp*PSDH expression level. We transformed *E. coli* BL21 for heterologous expression of either *Rp*PSDH or a SUMO-tagged variant (s*-Rp*PSDH), with the goal of supplementing β-hydroxylated nsAA and forming non-hydroxylated nsAAs through the action of native aminotransferases. We supplied 5 mM L-*threo*-phenylserine and monitored the formation of phenylalanine. We observed nearly 2-fold higher levels of phenylalanine formed when using the cells that express s-*Rp*PSDH, likely due to its higher expression level.

**Figure 3.**
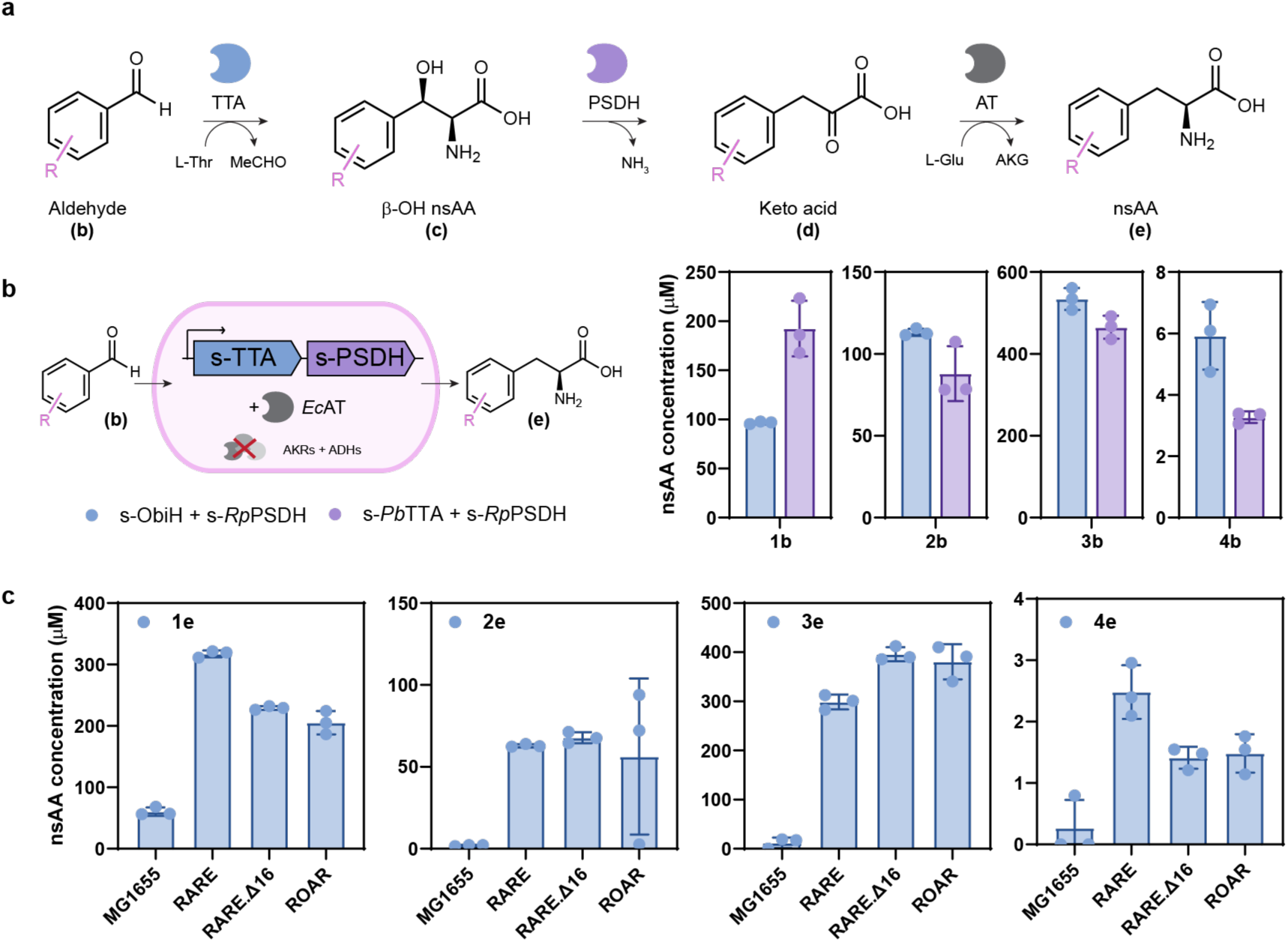
Demonstration of biosynthesis of encodable phenylalanine derivatives from aldehydes supplemented to live cells. (A) Designed biochemical pathway featuring reactions catalyzed by an L-TTA, PSDH, and AT. (B) Concentration of nsAA formed by the supplementation of 1 mM of each of the model aldehydes **1b**-**4b** to cells that are transformed to express a SUMO-tagged L-TTA (either s-ObiH or s-*Pb*TTA) and s-*Rp*PSDH, with a reliance on native expression levels of endogenous aminotransferases to form the desired nsAA products **1e**-**4e**. (C) Concentration of nsAA formed by the supplementation of 1 mM of each of the aldehydes **1b-4b** to four different strains (MG1655, RARE, RARE.Δ16, and ROAR) transformed to express s-*Pb*TTA and s-*Rp*PSDH.

We next sought to evaluate whether we could convert supplemented aryl aldehydes to non-hydroxylated nsAAs in growing cell cultures. We coupled one of the two highest performing SUMO-tagged L-TTAs from our previous screen (either s-ObiH or s-*Pb*TTA) with s-*Rp*PSDH on a single pCola vector for heterologous expression. We then monitored the conversion of 1 mM aldehydes supplemented to live cells at mid-exponential growth phase (OD_600_ = 0.4-0.6) to the anticipated phenylalanine derivatives (**Fig. 3b**). We were excited to see desired nsAA products **1e-3e** in the hundred micromolar range when supplying **1b**-**3b** and that we still produced detectable **4e** when supplying **4b** despite the poor activity of all the L-TTAs in our collection on **4b**. While relatively low, the 5 µM titer of **4e** achieved with our worst performing chemistry may nonetheless remain applicable for genetic code expansion.^37^ To determine if native TyrB levels could be limiting reaction yields, we investigated TyrB overexpression as well as supplementation of excess amine donor (L-Glu). We co-transformed the pCola vector containing s-TTA and s-*Rp*PSDH with a pZE vector containing TyrB in the RARE.Δ16 strain and measured nsAA titers in the presence and absence of excess L-Glu 20 h after 1 mM aldehyde addition. However, we did not observe a benefit from TyrB overexpression or L-Glu supplementation under the conditions tested. Finally, to understand the influence of the aldehyde stabilizing host strains on biosynthesized nsAA titer, we transformed multiple *E. coli* strains with our pathway plasmid containing only s-TTA and s-*Rp*PSDH. We chose the wild-type *E. coli* MG1655 strain, the originally published RARE strain that contains 6 deletions of aldehyde reductases (RARE.Δ6), the RARE.Δ16 strain mentioned earlier, and the ROAR strain, which contains 6 deletions of aldehyde oxidases in addition to the deletions in RARE.^35^ We compared production of nsAAs **1e-4e** 20 h after IPTG induction and supplementation with the corresponding aldehyde precursor. Notably, we observed minimal titers of each of these four model products when using the MG1655 strain. Comparatively, we observed >5-fold titers of **1e** in RARE, >30-fold titers of **2e** in RARE and RARE.Δ16, >30-fold titers of **3e** in RARE.Δ16 and ROAR, and >9-fold titers of **4e** in RARE (**Fig. 3c**). These results highlighted the value of harnessing an aldehyde stabilizing chassis to allow the implementation of this designed biochemical pathway in live cells.

### Demonstration of the extended biosynthetic pathway starting from carboxylic acids

We next investigated whether we could extend our pathway to demonstrate biosynthesis of our model nsAAs **1e**-**4e** from their associated carboxylic acids **1a**-**4a**. Anticipating that activity and soluble protein expression of CAR homologs could vary, we tested a panel of of six CAR homologs from our previously published collection.^38^ These were individually cloned onto a pZE vector to evaluate activity in live cell contexts in the aldehyde-stabilizing RARE.Δ16 strain. The CARs were co-expressed with Sfp from *Bacillus subtilis (Bs*Sfp), a phosphopantetheinyl transferase required for CAR activation^39^. We supplied 2 mM of the carboxylic acids **1a**-**3a** or 1 mM of **4a** (due to lower solubility) to RARE.Δ16 cells induced to express six different CARs during mid-exponential phase growth (**Fig. 4a**). We monitored reactions for 2 h given an interest in identifying fast-acting CAR variants. We detected aldehyde products in every case and observed that aldehyde product titers were comparable in many instances. Many CARs demonstrated high activity on acids **1a-3a**, while all showed moderately low activity on **4a**. We proceeded with the *Mav*CAR construct for subsequent experiments as we had known the protein to express well under various conditions.

**Figure 4.**
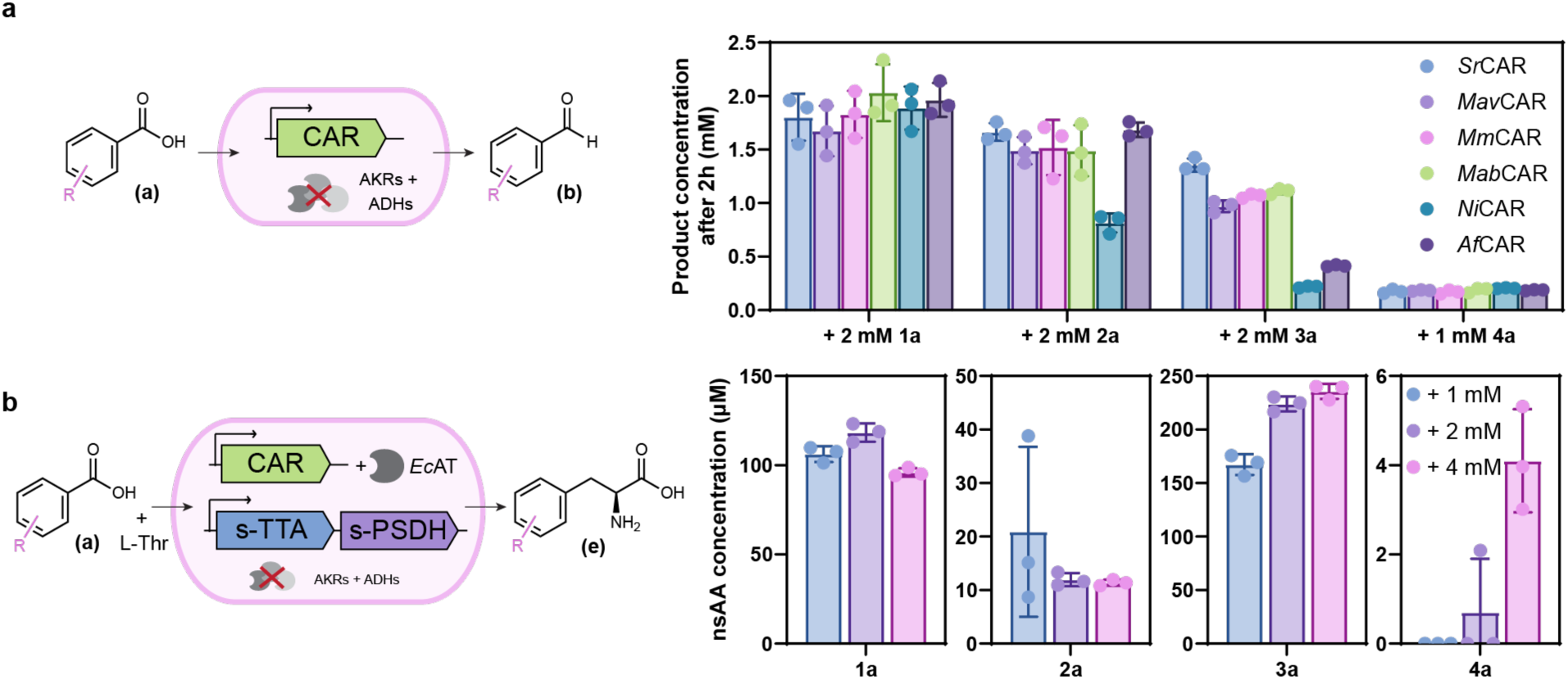
Demonstration of biosynthesis of encodable phenylalanine derivatives from carboxylic acids supplemented to live cells. (A) Comparison of the performance of cells that are transformed to express one of six carboxylic acid reductase (CAR) homologs for converting supplemented carboxylic acids **1a-4a** to their corresponding aldehydes. An endpoint of 2 h is chosen to identify fast-acting variants. (B) Concentration of nsAA formed by the supplementation of each of the model carboxylic acids to cells that are transformed to express *Mav*CAR, s-ObiH, and s-*Rp*PSDH, with a reliance on native expression levels of endogenous aminotransferases to form the desired nsAA products **1e**-**4e**. Three different concentrations of carboxylic acids are supplied for each chemistry as shown.

To determine whether co-expression of *Mav*CAR could allow a single strain to carry out the extended biosynthetic pathway from supplemented carboxylic acids, we transformed *E. coli* strains for heterologous expression of *Mav*CAR, *Bs*Sfp, either s-*Pb*TTA (for **1-3)** or s*-*ObiH (for **4**), and s-*Rp*PSDH. Initially, we utilized a two-plasmid system, both containing a medium-high (15-40) copy number origin of replication (ori) for both the *Mav*CAR with *Bs*Sfp (pZE vector, ColE1 ori) and the s-TTA with s-*Rp*PSDH (pCola vector, ColA ori) constructs. However, this initial design resulted in impacts on cell growth and poor titers. Consequenty, we moved the *Mav*CAR construct to a lower copy number plasmid (pACYC, p15A ori). With the second design, consisting of the original pCola vector containing the s-TTA and s-*Rp*PSDH co-transformed with the pACYC vector containing *Mav*CAR and *Bs*Sfp, we observed more robust growth. We then sought to understand the influence of exogenously supplied substrate concentration on desired nsAA product titers by supplying 1 mM, 2 mM, or 4 mM **1a**-**4a**. Endpoint HPLC measurements of the culture media indicated sucessful production of the desired nsAAs, with **4e** detectable in all replicates only when 4 mM **4a** was supplied (**Fig. 4b**). These results indicate that the influence of substrate concentration on product titers is chemistry-dependent.

### Identification of relevant systems for site-specific incorporation of targeted nsAAs

To achieve site-specific incorporation of the nsAAs biosynthesized by this pathway, we must identify a suitable genetically encoded orthogonal translation system (OTS) that would be capable of catalyzing aminoacylation of an orthogonal tRNA with the nsAA at biosynthetically attainable titers. To measure this, we must also co-express a reporter protein that contains a codon targeted for suppression by the OTS. We began by performing a well-established fluorescence assay with commercially sourced nsAAs **1e** and **4e** (0.5 mM) or no nsAA supplemented to engineered strains of *E. coli*. These *E. coli* strains were transformed to express one of five previously reported OTSs derived from the *Methanocaldococcus jannaschii* tyrosyl-tRNA synthetase (*Mj*TyrRS)/tRNA pair and a green fluorescent protein (GFP) reporter containing an in-frame UAG codon targeted for suppression (**Fig. 5a**). We performed an initial screen to identify the OTSs that maximized the fold-change increase in fluorescence normalized to optical density (FL/OD) between the presence and absence of each nsAA.

**Figure 5.**
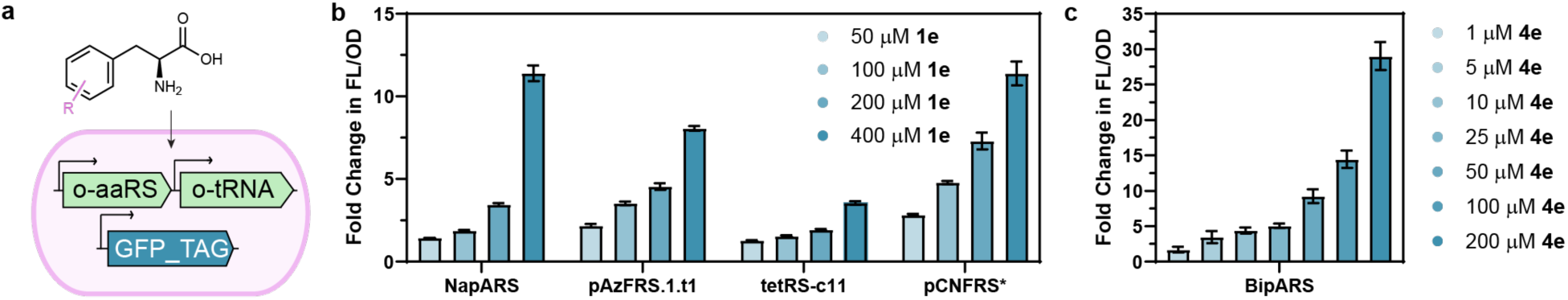
Aminoacyl-tRNA synthetase screening for nsAA incorporation at low externally supplemented concentrations. (A) Depiction of the chemistry and genetic constructs corresponding to the well-established fluorescence assay used for preliminary evaluation of relevant orthogonal translation systems (OTSs). (B) and (C) show measurement of dose-response (FL/OD) of the best identified OTS(s) for each chemistry at lower concentrations relevant to the titers achieved by biosynthesis for **1e** and **4e**, respectively.

To examine the dose-response of nsAA incorporation rates at concentration ranges more relevant to our biosynthetic titers, we selected the better performing OTSs for each chemistry from our initial screen. Fortunately, several OTSs for **1e** (**Fig. 5b**) and the BipARS for **4e** (**Fig. 5c**) exhibited ≥ 3.5-fold-change in FL/OD at concentrations as low as 100 µM **1e** or 5 µM **4e**, which correspond to nsAA titers attainable using our biosynthetic pathway. The high affinity of the BipARS for **4e** is a fortunate coincidence for our pathway, given its low micromolar yield of **4e** even when starting from the aldehyde **4b** or higher concentrations of the acid **4a**.

### Site-specific incorporation of biosynthesized nsAAs for control of protein translation using aldehydes or carboxylic acids

With suitable OTSs identified, we prepared genetic constructs that enabled co-expression of OTS, reporter protein that contains the in-frame UAG codon, and biosynthetic pathway, all in a single strain. We began with the core biosynthetic pathway starting from aldehydes with the aspiration of incorporating biosynthesized **4e** from supplemented 2 mM **4b** in LB media (**Fig. 6a**). We performed the biosynthesis and incorporation experiment in the *E. coli* RARE.Δ16 strain co-transformed with a vector (pCola) containing the biosynthetic pathway enzymes (s-ObiH and s-*Rp*PSDH) along with a vector (pRepSynth) containing a reporter protein (GFP_TAG) and OTS (BipARS/tRNA_CUA_ pair). After purification, trypsin digestion, and LC/MS/MS analysis of our reporter protein, we were delighted to see high-fidelity incorporation of biosynthesized **4e** (98%) when conducting a search that considered masses that could correspond to common misincorporation targets of Tyr, Trp, and Phe. This result illustrates that our combined system for biosynthesis and incorporation of phenylalanine derivatives can function robustly even for the model chemistry that was least tolerated by our biosynthetic pathway enzymes.

**Figure 6.**
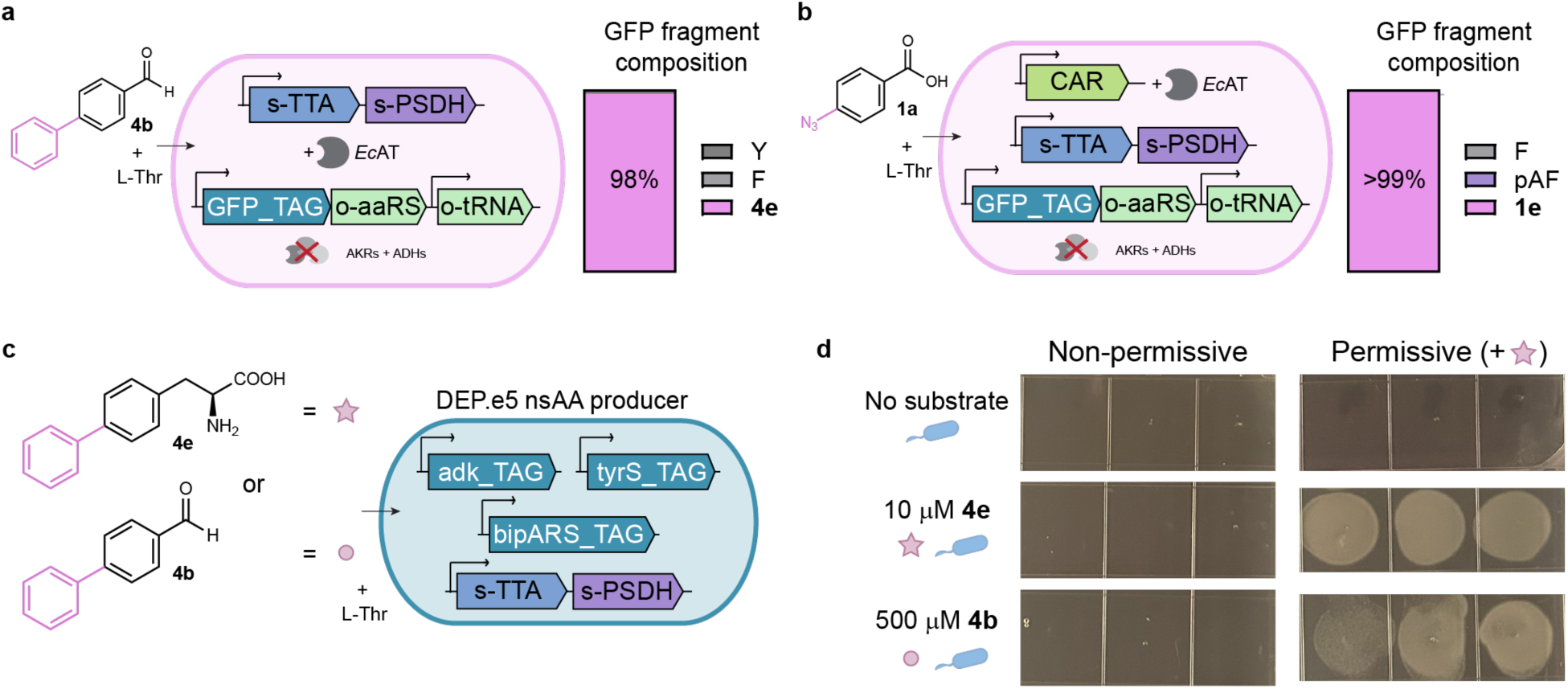
Site-specific incorporation of biosynthesized nsAAs from aldehyde or carboxylic acid precursors using a single engineered strain of *E. coli*. (A) Genetic construct design and digested peptide mass spectrometry result indicating high-fidelity site-specific incorporation of biphenylalanine (**4e**) within GFP reporter protein when supplying cells 2 mM biphenylaldehyde (**4b**) as a representative aldehyde, which corresponded to the lowest yield product made by the pathway. (B) Genetic construct design and digested peptide mass spectrometry indicating high-fidelity site-specific incorporation of 4-azido-phenylalanine (**1e**) within GFP reporter protein when supplying cells 2 mM 4-azido-benzoic acid (**1a**), which is far less expensive on a normalized basis than the nsAA. (C) Solid-media growth and escape from biocontainment assay indicating the successful extension of synthetic auxotrophy to include reliance on the aldehyde **4b**. This was done by transforming a previously engineered and evolved strain that requires **4e** (DEP.e5) for heterologous expression of our core biosynthetic pathway. The strain already contains a BipA-dependent BipARS and essential genes that are modified to contain in-frame UAG codons. Thus, transformation of this strain with our pathway does not result in observable escape under the conditions tested, and it does allow the strain to grow if precursor **4b** is provided.

To determine whether we could convert carboxylic acids to incorporated nsAAs, we selected **1e** as a commercially relevant target given the significantly lower cost of **1a** compared to **1e**. We transformed cells of the ROAR strain^40^ for heterologous expression of the biosynthetic pathway, the OTS, and the reporter and supplied 2 mM of **1a** precursor (**Fig. 6b**) to live cells in LB media. We utilized a three-plasmid system consisting of 1) a vector (pZE) containing *Mav*CAR and *Bs*Sfp, 2) a vector (pCola) containing the biosynthetic pathway enzymes (s-ObiH and s-*Rp*PSDH) and 3) a vector (pRepSynth) containing a reporter protein (GFP_TAG) and OTS (NapARS/tRNA_CUA_ pair). After purification, trypsin digestion, and LC/MS/MS analysis of our reporter protein, we observed high-fidelity incorporation of biosynthesized **1e** (>99%), with minor azide reduction to 4-amino-phenylalanine (pAF) observed (0.3%). This approach could offer substantial cost savings for commercial-scale manufacturing of therapeutic proteins that contain azide functionality.

### Control of bacterial survival by provision of biphenylaldehyde

The ability to couple biosynthesis and incorporation of nsAAs through semi-synthesis strategies that harness low-cost synthetic precursors creates new opportunities in synthetic biology and biosensing where cellular function can now be governed by the concentration of certain aryl aldehydes or carboxylic acids. An exemplary cellular function is survival, where previous work has demonstrated the ability of nsAAs to control survival by mediating the full-length translation and proper folding of essential proteins. Pioneering efforts in this direction reported synthetic auxotrophs, which are intrinsically biologically contained due to their reliance on a chemical not found in nature.^41,42^ *E. coli* strains that were designed to rely on **4e** included some that exhibit no observable escape under various conditions, including after 100 days of continuous culturing.^37^ Importantly for the present effort, continuous culturing resulted in adaptation to the applied pressure of low micromolar concentrations of supplemented **4e**. As chiral molecules, **4e** and other nsAAs may be cost-prohibitive to supply at scales required for environmental applications of biocontained strains.

To create a synthetic auxotroph responsive to biphenylaldehyde (**4b**), we transformed an evolved biphenylalanine-dependent strain, DEP.e5, for heterologous expression of our core biosynthetic pathway, consisting of s-ObiH and s-*Rp*PSDH (**Fig. 6c**). We then performed a growth-based assay that measures the ability of an alternative molecule than biphenylalanine (**4e**) to support growth in liquid media. We were interested in measuring growth in the medium intended to be permissive as well as potential escape from biocontainment in the medium lacking a synthetic molecule. While the DEP.e5 strain has not been observed to escape, it is possible that its transformation with such a promiscuous pathway could enable it to biosynthesize unintended nsAAs from culture media components or native metabolites. We include a no substrate case as a negative control, a case of supplying 10 µM **4e** as a positive control, and our experimental case of supplying 500 µM **4b** (**Fig. 6d**). Cells of the DEP.e5 strain transformed with our pathway were only able to grow when **4e** or ≥ 250 µM **4b** were provided in liquid media. We subsequently confirmed growth was due to reliance on the supplemented synthetic molecules by plating on solid media permissive (containing **4e**) or non-permissive (without **4e**) conditions and observed growth only under permissive conditions with cells previously grown in the presence of **4e** or **4b**. Collectively, these results indicate the ability to use our pathway to create synthetic auxotrophs that can grow in the presence of user-defined synthetic aldehydes.

## Discussion

The current paradigm of nsAA synthesis, isolation, and supplementation to live cells cultures for site-specific incorporation within proteins is a major constraint on the application of genetic code expansion at industrially relevant scales and for emerging environmental uses. Here, we demonstrate the design of a synthetic biochemical pathway that functions in live bacterial cells and its coupling to orthogonal translation systems for combined nsAA biosynthesis and incorporation within a single strain. This approach reduces the problem of sourcing a phenylalanine derivative to a problem of sourcing an aryl aldehyde or aryl carboxylic acid, which are simple achiral building blocks. These synthetic compounds are widely available, and because carboxylic acids are also ubiquitous in metabolism, this creates opportunities to biosynthesize the precursors towards total biosynthesis from renewable feedstocks. This coupling of metabolic engineering to genetic code expansion could address other bottlenecks in genetic code expansion, such as transport, nsAA toxicity (by maintaining a lower steady state in some cases), or nsAA concentration (by increasing intracellular concentration in other cases).

The coupling of nsAA biosynthesis and incorporation was accomplished using a pathway consisting of up to five highly-polyspecific enzymes that are capable of functioning together in a single cell. Given the promiscuity of the biosynthetic pathway observed *in vitro*,^26^ we expect that the coupled system in live cells will accept additional aryl aldehyde and carboxylic acid substrates. Our system demonstrated that even at low biosynthetic yields for certain chemistries, coupling with a high affinity and high-fidelity orthogonal translation system can still achieve protein incorporation with high fidelity. Further optimization of expression levels, enzyme activity, and substrate tolerance, particularly for bulky substrates, and even protein engineering of our naturally occurring pathway enzymes may enhance the biosynthetic platform. Alternatively, the engineering of OTSs to exhibit higher affinity towards nsAAs has been relatively underexplored, though there has been recent successful evolution campaigns that characterize lower limit of detection.^43^ Engineering higher affinity OTSs should serve as a less burdensome means of improving target protein yield when coupling nsAA biosynthesis and incorporation. By concluding this study with the creation of a strain whose survival depends on the biosynthesis and incorporation of biphenylalanine from biphenylaldehyde, we now have a unique selection platform for directed evolution or adaptive laboratory evolution of any of the steps in this process.

## Acknowledgments

We sincerely thank Y. Yu and P.N. Asare-Okai in the Mass Spectrometry Facility at the University of Delaware for the mass spectrometry analysis and assistance that is supported by the National Institute of General Medical Sciences or the National Institutes of Health under award P20GM104316. We acknowledge support from the following funding sources: the Office of Naval Research Award No. N000142212536 (to A.M.K.); Department of Energy Joint Genome Institute under proposal 506446, a DOE Office of Science User Facility, supported by the Office of Science of the U.S. Department of Energy operated under contract no. DE-AC02-05CH11231 (to A. M. K); the National Institutes of Health Centers of Biomedical Research Excellence under grant 5P20GM104316-08 (to A. M. K.); the National Science Foundation Division of Chemical, Bioengineering, Environmental, and Transport Systems Award No. CBET-2032243 (to A.M.K. and A.P.S); and the National Institute of General Medical Sciences of the National Institutes of Health under a Chemistry-Biology Interface Training Grant T32GM133395 (to S.R.A. and M.A.J.). Additionally, we would like to thank the members of the Kunjapur group for thoughtful discussions towards this manuscript.

